# *SdiA*, a Quorum Sensing Transcriptional Regulator, Enhanced the Drug Resistance of *Cronobacter sakazakii* and Suppressed its Motility, Adhesion and Biofilm Formation

**DOI:** 10.1101/2022.02.02.478922

**Authors:** Chuansong Cheng, Xiaotong Yan, Binxiong Liu, Tao Jiang, Ziwen Zhou, Dongwei Zhang, Huayan Wang, Dengyuan Chen, Changcheng Li, Ting Fanga

## Abstract

*Cronobacter sakazakii* is a common foodborne pathogen, and the mortality rate of its infection is as high as 40-80%. Quorum sensing is a regulation system of bacterial density-dependent multigene expression and is an important regulatory mechanism involved in adhesion, biofilm formation and virulence. *C. sakazakii* contains a QS signal molecular receiver, which is the LuxR receptor homolog *SdiA*, but its regulatory mechanism in *C. sakazakii* QS has not been defined. Here, we further determined the effect of *SdiA* on the QS system of *C. sakazakii*. The *SdiA* gene in *C. sakazakii* was knocked out by gene editing technology, and the biological characteristics of the Δ*sdiA* gene deletion strain of *C. sakazakii* were studied, followed by transcriptome analysis to elucidate its effects. The results suggested that *SdiA* enhanced the drug resistance of *C. sakazakii* but diminished its motility, adhesion and biofilm formation ability and had no effect on its growth. Transcriptome analysis showed that the deletion of the *SdiA* gene upregulated the expression levels of D-galactose operon genes (including *dgoR*, *dgoK*, *dgoA*, *dgoD* and *dgoT*) and flagella-related genes (*FliA* and *FliC*) in *C. sakazakii* and downregulated the expression levels of related genes in the type VI secretion system (*VasK* gene was downregulated by 1.53-fold) and ABC transport system (downregulated by 1.5-fold), indicating that *SdiA* was related to the physiological metabolism of *C. sakazakii*. The results of this study may be useful for clarifying the pathogenic mechanism of *C. sakazakii* and provide a theoretical basis for controlling bacterial infection.

**IMPORTANCE:** *Cronobacter sakazakii*, as an emerging opportunistic foodborne pathogen, was associated with sepsis, meningitis and necrotizing enterocolitis in neonates and infants, with a mortality rate of 40-80%. Quorum sensing plays an important regulatory role in the pathogenicity of *C. sakazakii*. Nevertheless, the regulatory mechanism of QS in *C. sakazakii* remains unknown. Here, we studied the QS transcriptional regulator *SdiA* of *C. sakazakii*. We revealed the regulatory mechanisms of *SdiA* in *C. sakazakii* cell adhesion, motility, biofilm formation and drug resistance. It was helpful to further explore the function of the *SdiA* gene, revealing the pathogenic mechanism of *C. sakazakii*. It will also provide a new target for therapeutic interventions targeting the pathogenicity of *C. sakazakii* and developing quorum-sensing inhibitors.

## INTRODUCTION

*Cronobacter sakazakii*, a spiral-shaped flagellated Gram-negative rod-shaped bacilli bacterium, is an emerging food-borne opportunistic pathogen that is primarily parasitic in human and animal intestines (1, 2). *C. sakazakii* comes from a wide range of sources, and Powdered Infant Formula (PIF) is considered the primary source of infection and medium for transmission (3–5). In addition, *C. sakakazaki* was also isolated from cheese, grains, fruits, vegetables, herbs and meats (6, 7). Neonates and young infants are susceptible to the bacterium that could cause severe clinical presentations of septicaemia, meningitis and necrotizing enterocolitis and produce possible life-threatening chronic neurologic sequelae (8, 9). Unfortunately, the death rate is as high as 40-80% (10).

Quorum sensing (QS) is a collaborative regulation system of bacterial density dependent on multigene expression, an environmental signal sensing system for bacteria to monitor their own population density, an important regulatory mechanism involved in adhesion, biofilm formation, virulence and so on (11–14). In most Gram-negative bacteria, two major QS systems are autoinducer-1 (AI-1) and autoinducer-2 (AI-2). The AI-1-type QS system is mainly responsible for communication between Gram-negative bacteria. LuxI and LuxR are two key proteins: LuxI synthase produces *N*-acyl homoserine lactones (AHLs) as its own inducers, and the LuxR transcription factor is their homologous receptor (15–19). The AI-2-type QS system is present in both Gram-negative and Gram-positive bacteria, and LuxS is a key protein (20, 21).

Interestingly, some Gram-negative bacteria encode LuxR receptors but do not produce AHLs because they lack LuxI synthase. For instance, *Klebsiella pneumoniae*, *Salmonella* and *Enterobacterium* do not have the LuxI synthase gene, and they do not produce AHL signaling molecules (11, 19). However, there is a chromosomal LuxR homolog called *SdiA* of these bacteria; these bacteria can use the *SdiA* sensor to sense the signaling molecules produced by other bacteria in the environment and then regulate the expression of their own related genes (22).

*SdiA* was identified as a “suppressor of cell division suppressors” that controls the transcription of *ftsQAZ* operons involved in cell division. It has been noted that in *enterohemorrhagic Escherichia coli* (EHEC) and *Enterobacteriaceae*, *SdiA* is involved in the regulation of many virulence factors, such as pili production, biofilm formation, adhesion and motility of cells (22–24). In addition,, *Salmonella* utilizes *SdiA* proteins to detect AHLs synthesized by other species and enhances *srgE* and *RCK* operon expression (20, 11). Recent studies have reported that *SdiA* has an effect on the survival of *C. sakazakii* under different environmental stresses, with improvement of *C. sakazakii* tolerance to heat, desiccation, osmotic and acid stress conditions (25). Nevertheless, the function of the QS receptor *SdiA* in *C. sakazakii* and its underlying mechanisms remain largely unknown. More studies are required to elucidate the exact functions of *SdiA* on the physiology of these microorganisms since a great level of complexity is observed as revealed by these previous works.

In this study, *gene editing* technology was used for the construction of an *SdiA* deletion mutant to investigate the role of *SdiA* in *C. sakazakii* pathogenicity by assessing the growth, biofilm formation, motility, adhesion and multidrug resistance of the QS receptor *SdiA* in *C. sakazakii.* Meanwhile, transcriptome analysis was conducted to elucidate the expression of *SdiA*-related genes.

## RESULTS

### Construction of a *C. sakazakii* CICC 21550 mutant strain

In this study, we constructed a *C. sakazakii* CICC 21550 strain in which the *SdiA* gene was deleted by gene replacement. Twenty-four single colonies were randomly selected from TSA containing chloramphenicol (17 μg/ml) agar plates for PCR analysis to screen the *SdiA* gene knockout strain. At the 5’ endpoint of the target fragment, a 1368 bp fragment was generated by pairing *SdiA*-outF and CmSeqF2 primers. At the 3’ endpoint, CmSeqR2 and *SdiA*-outR primer pairs produced a 1391 bp fragment. When both fragments were present, the chloramphenicol resistance gene fragment replaceed the *SdiA* gene fragment. The results indicated that both sides of clones 9 and 14 were amplified with strong specific PCR products in accordance with the expected length (Fig. 1D). After the chloramphenicol resistance gene was deleted, the length of the amplified product of the lateral primer was shortened to 1802 bp (Fig. 1B and C). This assay confirmed that we successfully constructed the *SdiA* deletion strain of *C. sakazakii* CICC 21550.

**Fig. 1.**
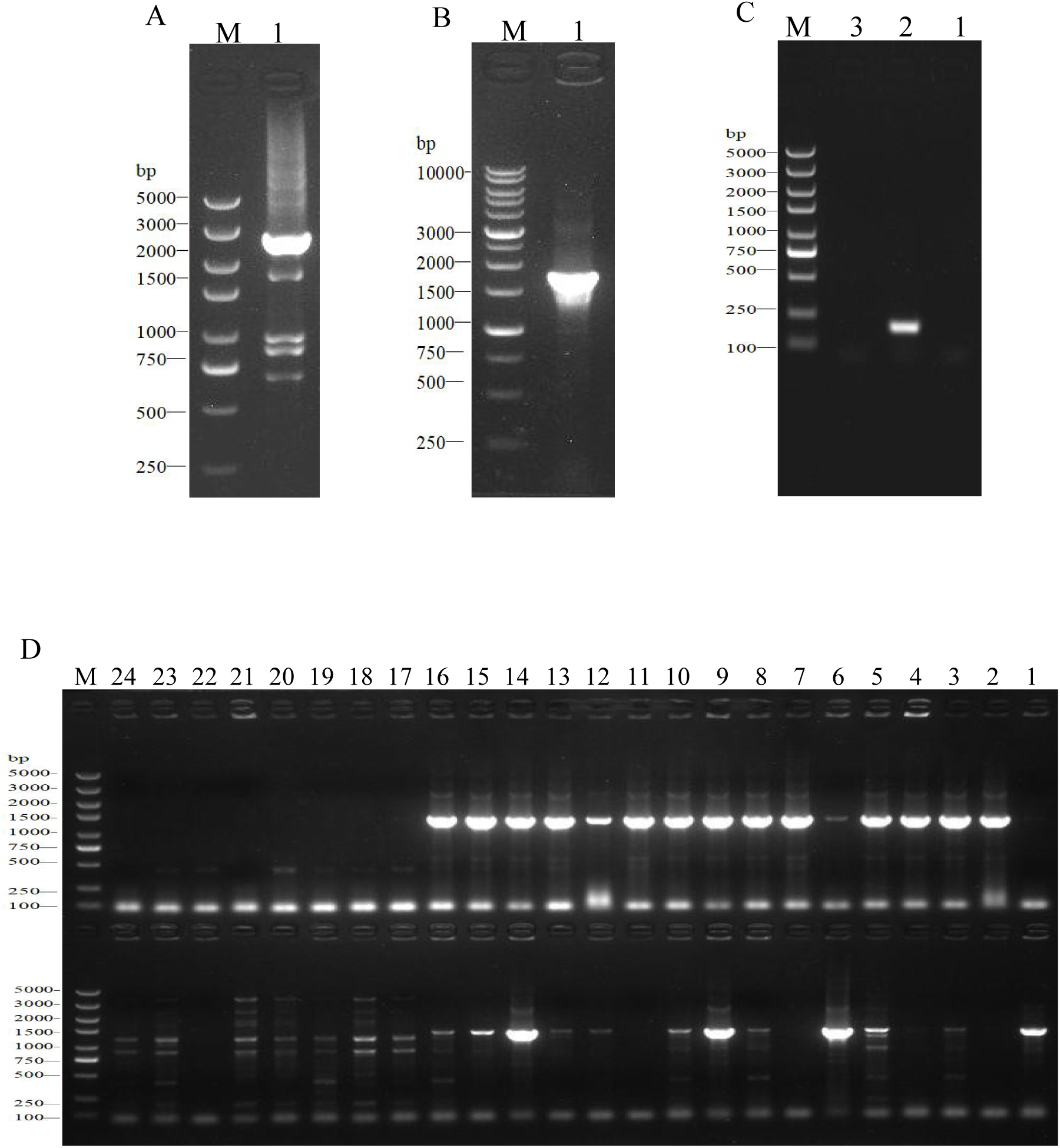
Construction and identification of the *SdiA* deletion mutant of *C. sakazakii* by gel electrophoresis. **(A)** Construction of *SdiA* gene target shooting fragment. M: DL5000 DNA marker; 1: *SdiA* gene target shooting fragment Δ*sdiA*::Cm (2650 bp). **(B, C)** Chloramphenicol resistance gene removal. **(B)** M: DL10000 DNA Marker; 1: Results of identification of lateral primers for colonies growing on non-resistant a TSA plates. **(C)** M: DL5000 DNA marker; 1: Identification results of lateral primers for colonies growing on non-resistant a LB plates; 2: Internal primer amplification results of the original strain (193 bp); 3: No template negative control amplification results. **(D)** Screening of *SdiA* gene knockout strain. M: DL5000 DNA marker; 1-24: Amplification results of primer pairs on both sides of 24 single colonies (Amplification results on both sides of 5’ endpoints above; Amplification results on both sides of the 3’ endpoint are shown below).

### Growth of *C. sakazakii* WT and Δ*sdiA* mutant strains in PIF

The growth model of the WT strain and Δs*diA* mutant strains at fluctuating temperatures was constructed using a one-step data analysis method (26). Fig. 2D shows the growth rate distribution of the Δ*sdiA* mutant and WT strains at 4-50℃, and their growth parameters were very similar. There were no significant differences, in that the absence of the *SdiA* gene had no effect on the growth of *C. sakakazaki*. Therefore, we combined the data of the WT and Δ*sdiA* mutant strains to analyze their growth (a total of 260 data points) in the recovered PIF at fluctuating temperatures of 8.4-48℃ based on the no lag-Cardinal model and Baranyi-Cardinal models of the one-step data analysis method (27, 28). The statistical results and estimated values of each parameter are shown in Table 2 and Table 3. The observed values and model prediction curves of the WT and Δ*sdiA* mutant strains at each temperature are shown in Fig. 2A, B and C. Fig. 2E shows the effect of temperature on the growth rate of the WT and Δ*sdiA* mutant strains. The results showed that no lag-Cardinal model and Baranyi-Cardinal model had the same fitting effect on the growth of the *C. sakazakii* WT and Δ*sdiA* mutant strains.

**Fig. 2.**
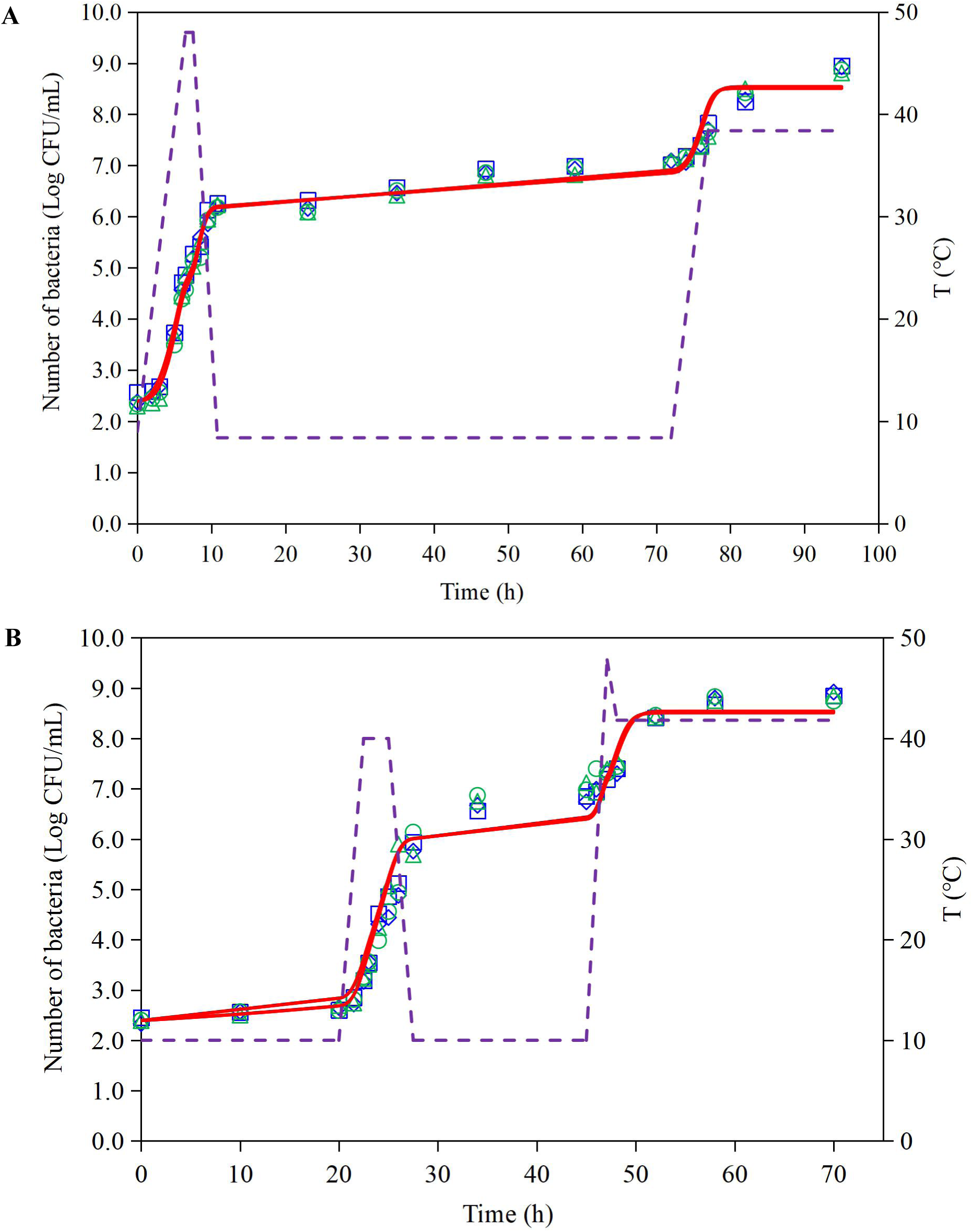
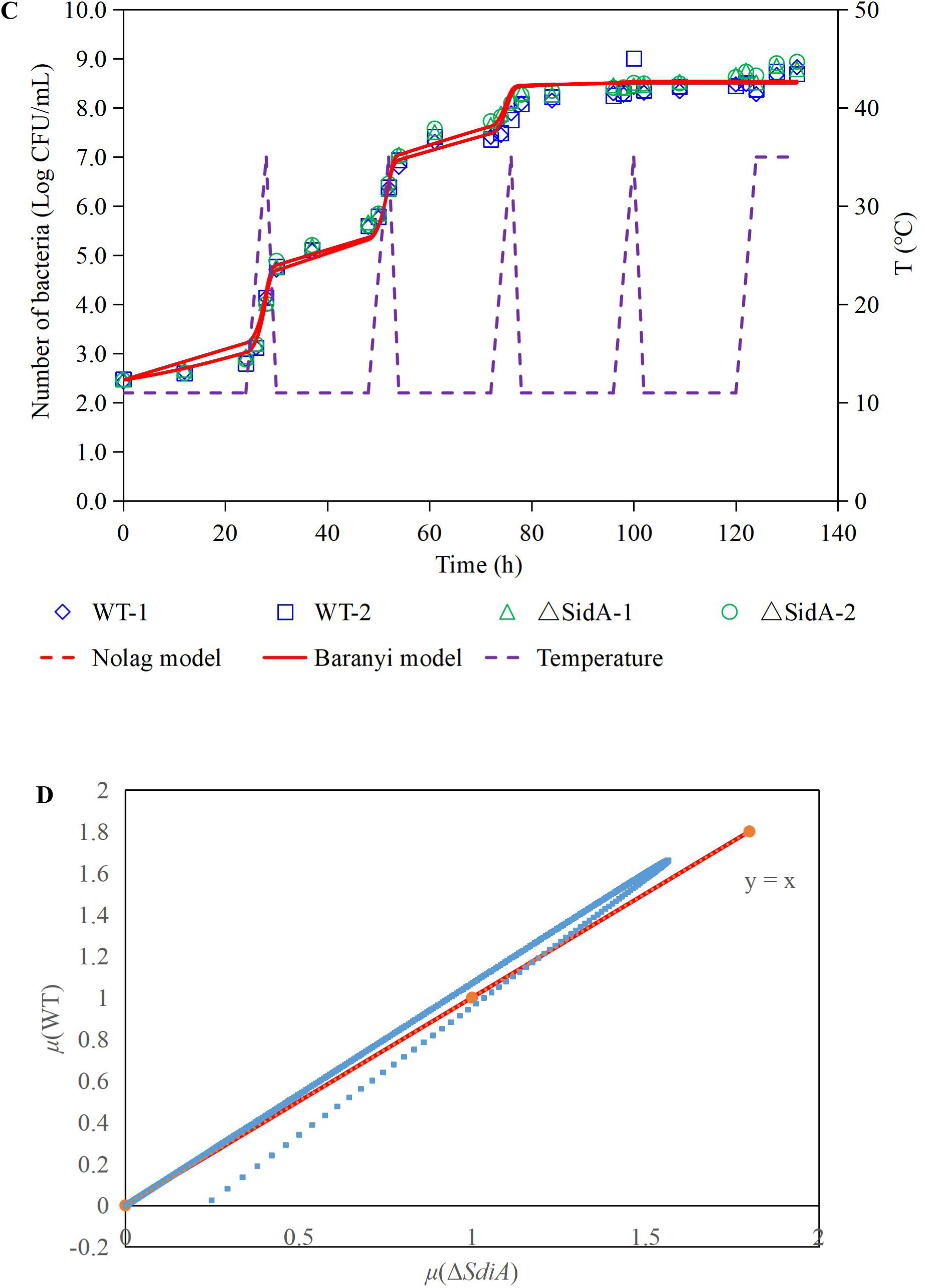
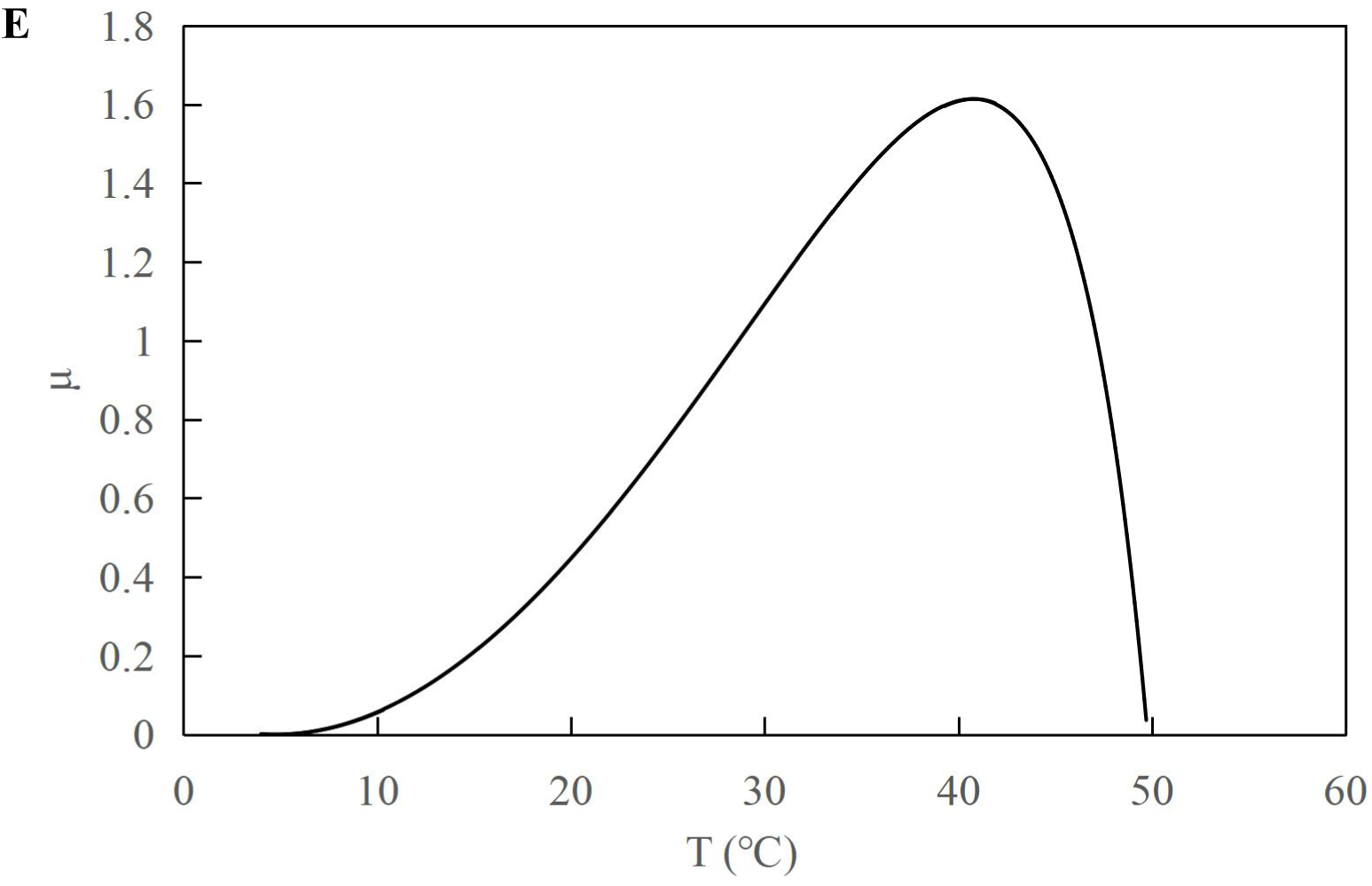
One-step analysis for *C. sakazakii* wild-type and Δ*sdiA* mutant strains. **(A, B, C)** One-step fitting analysis of the growth curve of *C. sakazakii* WT and Δ*sdiA* mutant strains in powdered infant formula recovery. **(D)** Growth rate distribution of Δ*sdiA* mutant and WT strains. **(E)** One-step analysis: Effect of temperature on growth rate of WT and Δ*sdiA* mutant strains.

**Table 2.**
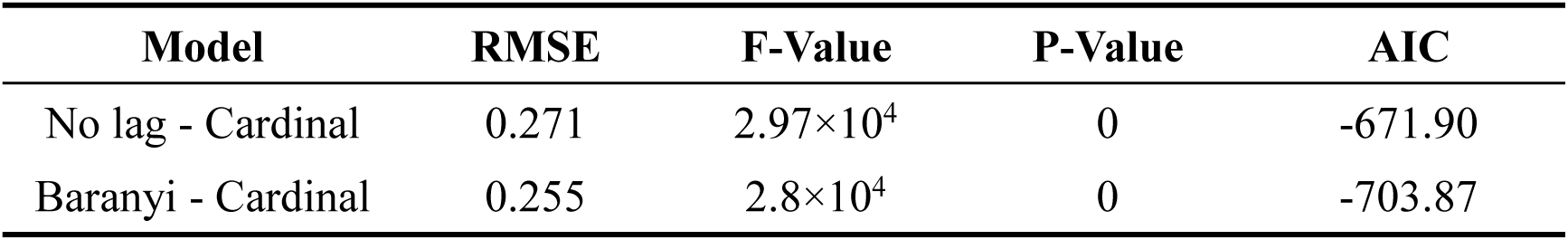
One-step analysis results of WT and Δ*sdiA* mutant strains.

**Table 3.**
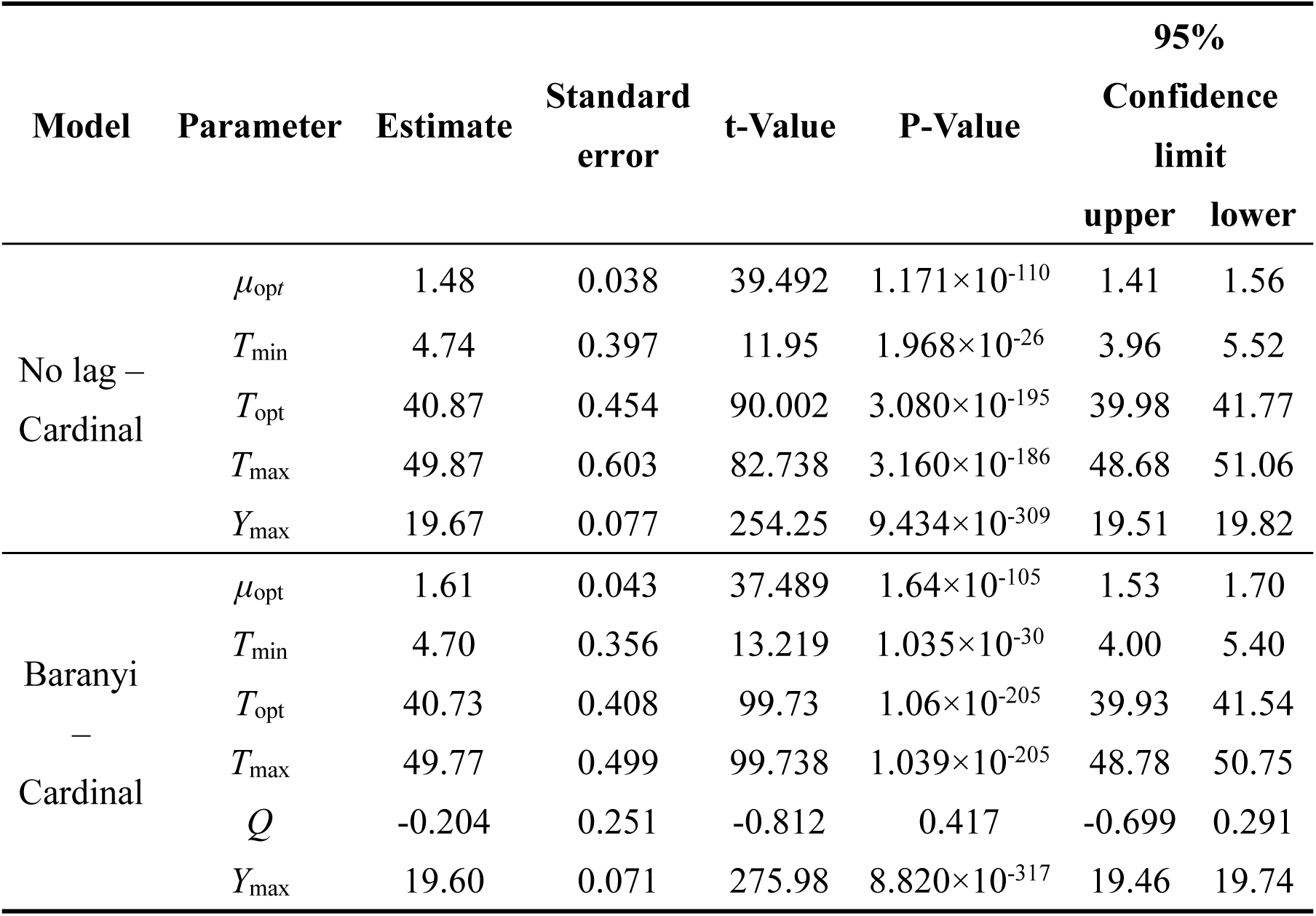
Parameter estimation of no lag - Cardinal and Baranyi - Cardinal models of WT and Δ*sdiA* mutant strains.

Hence, we further calculated the influence of *SdiA* gene deletion on the growth parameters of *C. sakazakii* and the growth changes of *C. sakazakii* in PIF. A mathematical model for the growth of *C. sakazakii* WT and Δ*sdiA* mutant strains in the recovered PIF was established by a one-step method. In this study, *C. sakazakii* WT and Δ*sdiA* mutant strain parameters were combined to be estimated by the no lag-Cardinal model and Baranyi-Cardinal model; the minimum growth temperatures estimated by the two models were 4.74℃ and 4.70℃, respectively, and the highest growth temperatures were 49.87℃ and 49.77℃, respectively. The optimum growth temperatures were 40.87℃ and 40.73℃, respectively, and the optimum growth rates were 1.48 h^-1^ and 1.62 h^-1^, respectively. In addition, the maximum growth concentrations estimated by the no lag-Cardinal model and Baranyi-Cardinal model were 19.67 Ln CFU/ml (8.54 log CFU/ml) and 19.60 Ln CFU/ml (8.51 log CFU/ml), respectively.

### Lack of *SdiA* increases the swarming and swimming motility of *C. sakazakii*

According to Fig. 3A, after deletion of the *SdiA* gene in *C. sakazakii*, the swimming motility of the mutant strain (Δ*sdiA*) was markedly enhanced. Nonetheless, the WT strain barely spread after 21 h of culture on TSA plates with 0.3% agar. The Δ*sdiA* mutant strain showed a trend of spreading after 6 h of culture. After incubation for 21 h, the Δ*sdiA* mutant strain spread to the entire plate, and the swimming motility of the Δ*sdiA* mutant strain showed a linear increase (Fig. 3C). Furthermore, the Δ*sdiA* mutant strain showed significantly higher swarming motility than the WT strain and rapidly spread after 18 h of culture, showing an exponential growth trend (Fig. 3A and D). Although the WT strain indicated a tiny trend of outward diffusion, their movement was still not visible after 36 h of culture. In the transcriptomics study of *C. sakazakii* WT and Δ*sdiA* mutant strains, we found that the expression levels of *FliA* and *FliC-*related genes regulating the flagellate of *C. sakazakii* were significantly increased (Tables S1, S3 and S4), which resulted in enhanced flagellate activity and significantly enhanced swarming and swimming motility of *C. sakazakii*.

**Fig. 3.**
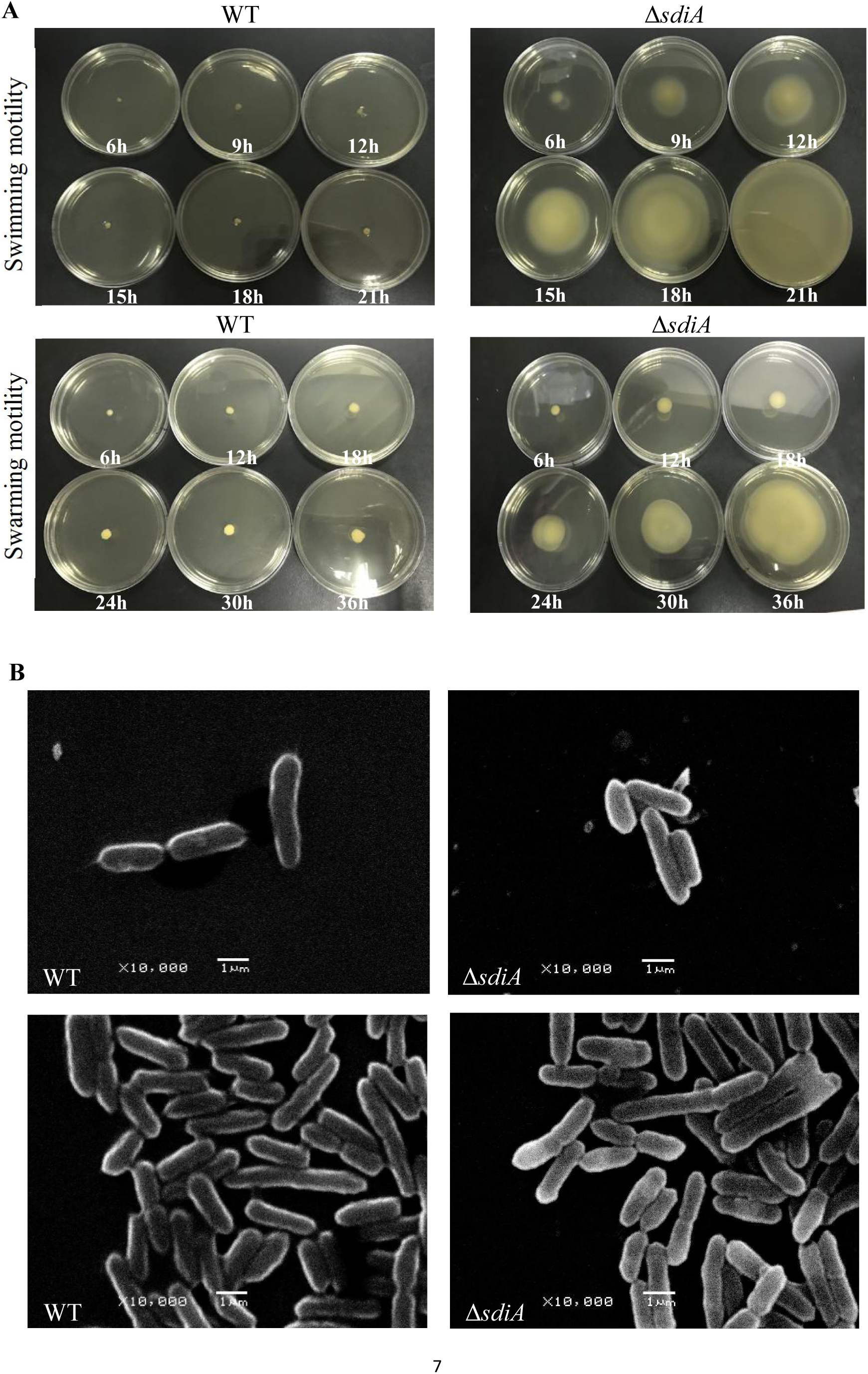
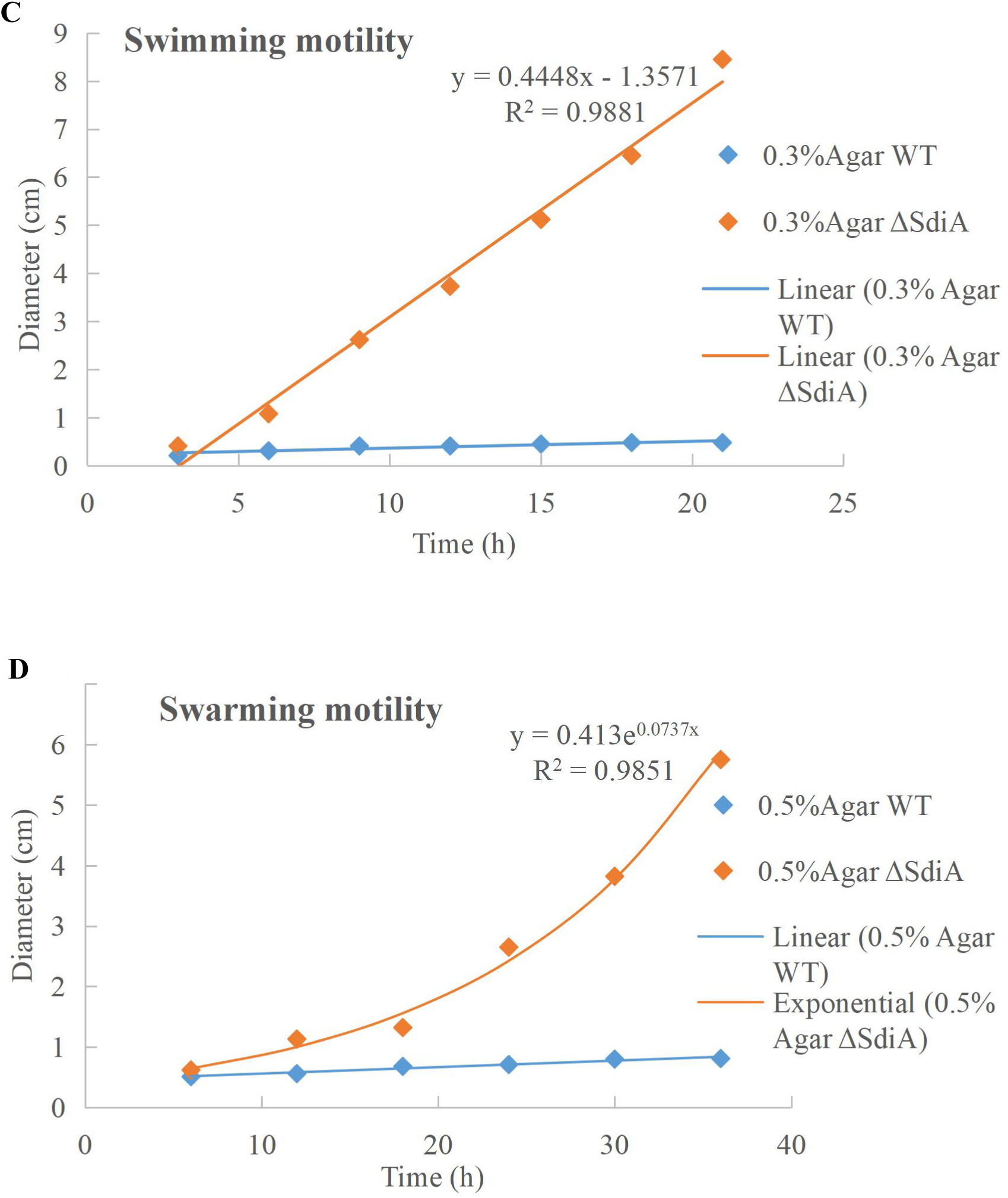
*C. sakazakii* WT and Δ*sdiA* mutant strains cell viability and morphology differences. **(A)** WT and Δ*sdiA* mutant strains activity assay. **(B)** WT and Δ*sdiA* mutant strains scanning electron microscopy (SEM) images. **(C, D)** WT and Δ*sdiA* mutant strains activity assay curve.

### Lack of *SdiA* increases the biofilm formation and surface adhesion capabilities of *C. sakazakii*

*SdiA* acts as a biofilm regulator in many foodborne pathogens, but its role in *C. sakazakii* remains unclear. Thus, to evaluate whether *SdiA* was also involved in the biofilm formation of *C. sakazakii*, we compared the biofilm formation ability of WT and Δ*sdiA* mutant strains. Compared with the WT strain of *C. sakazakii*, the amount of biofilm formation of the *SdiA* gene deletion strain increased by 38.54%, and the Δ*sdiA* mutant strain showed more biofilm formation (Fig. 4A).

**Fig. 4.**
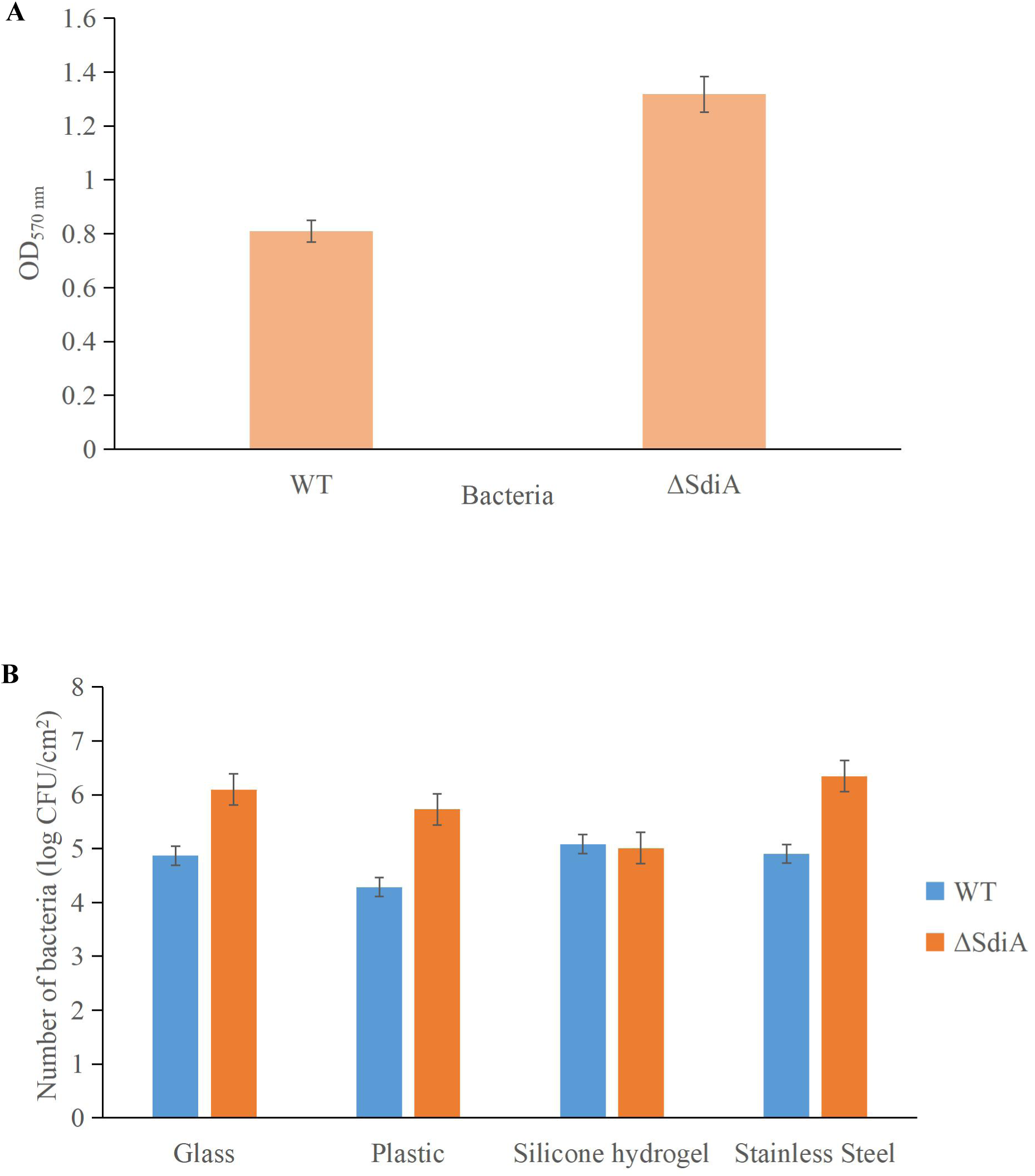
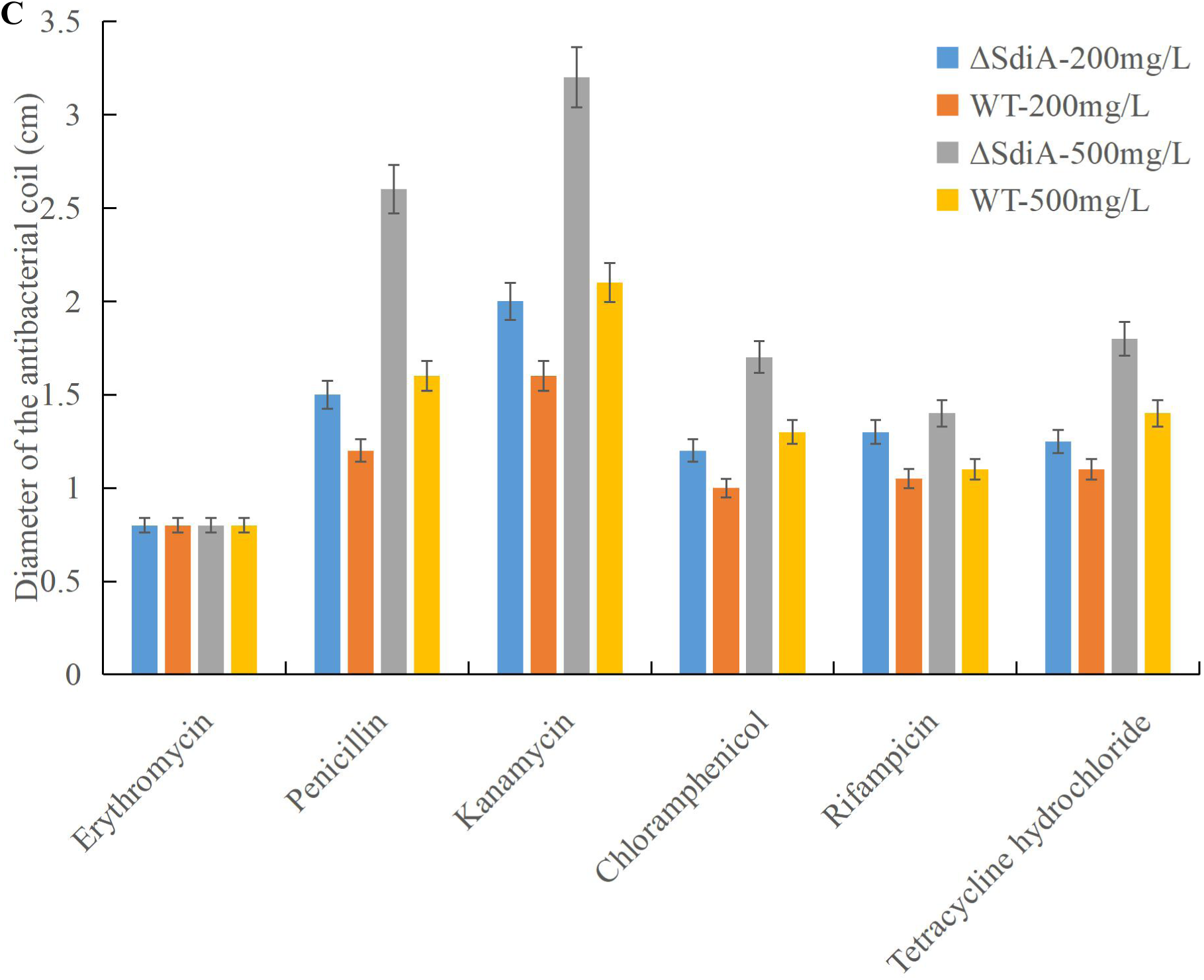
*C. sakazakii* WT and Δ*sdiA* mutant strains biofilm formation, surface adhesion capabilities and antibiotic resistance assay. **(A)** Optical density values of CV solutions obtained from biofilms of *C. sakazakii* WT and Δ*sdiA* mutant strains grown in the TSB at 37°C. (**B**) Number of adhered *C. sakazakii* WT and Δ*sdiA* mutant strains to different substratum. **(C)** *C. sakazakii* WT and Δ*sdiA* mutant strains antibiotic resistance assay. All experimental datas were plotted as mean ± standard from triplicate determinations and statistical analysis was performed with SPSS 17.0 Significance of WT and Δ*sdiA* mutant strains were established through t test using P values < 0.05.

We found that the *SdiA* gene regulates the adhesion of *C. sakazakii* to stainless steel, glass and plastics. The adhesion of the WT strain to glass, plastic, silicone hydrogel and stainless steel surfaces was 4.87, 4.29, 5.08 and 4.90 log CFU/cm^2^, respectively, at 30℃ for 15 min. The adhesion of the Δ*sdiA* mutant strain to glass, plastic, silicone hydrogel and stainless steel was 6.10, 5.73, 5.01 and 6.35 log CFU/cm^2^, respectively (Fig. 4B). The *SdiA* deletion resulted in increased adhesion of *C. Sakazakii* on glass, plastic and stainless steel (P<0.05), but there was no significant difference in adhesion between the WT and Δ*sdiA* mutant strains on the silica hydrogel surface. The adhesion of *C. sakazakii* was affected by the dielectric surface, and the adhesion of WT on the plastic surface was lower than that of the other three dielectric materials. However, the adhesion of the Δ*sdiA* mutant strain on the silica hydrogel surface was lower than that of the other three media materials. Through our transcriptomic studies, we also found that the expression of the ATP-active membrane-associated protein *VasK* in the VI secretion system, which is important to the regulation of *C. sakazakii* adhesion, was downregulated by 1.53-fold in *SdiA* mutant strains (Tables S2, S3 and S4). PIF packaging, brewing and drinking bottles were mainly made of stainless steel, glass, plastic and silicone hydrogel materials, and *C. sakazakii* was the main food-borne pathogen in PIF. By studying the adhesion of the *SdiA* mutant and WT strains to these materials, we can make it clear that the key to controlling the contamination of *C. sakazakii* during the processing and brewing of PIF was the targeted regulation of the *SdiA* gene.

### Lack of *SdiA* decreases the multidrug resistance of *C. sakazakii*

As shown in Fig. 4C, when the *SdiA* gene was deleted, antibiotic resistance of *C. sakazakii* showed a general decrease. With the increase in antibiotic concentration, the antibacterial zone of antibiotics showing antibacterial effect was larger, among which kanamycin showed the most obvious antibacterial effect and showed relatively good antibacterial effect on WT and Δ*sdiA* mutant strains. Second, penicillin has a certain bacteriostatic effect, but compared with kanamycin, it does not show a strong bacteriostatic effect. Chloramphenicol, rifampicin and tetracycline hydrochloride exhibited approximately the same antibacterial effect, with a slight antibacterial effect. Among the 6 antibiotics, erythromycin showed poor antibacterial activity against *C. sakazakii*, and neither WT nor Δ*sdiA* mutant strains showed an antibacterial zone, showing no visible antibacterial effect. By studying the changes in the transcript level of the *C. sakazakii SdiA* mutant and the wild-type strains, we found that the expression level of related genes in the ABC transport system regulating drug resistance of *C. sakazakii* was downregulated by 1.5-fold (Tables S2, S3 and S4) (29, 30), which revealed the mechanism by which the *SdiA* gene enhances drug resistance in *C. sakazakii*.

### Lack of *SdiA* alters the expression of genes involved in motility, the virulence factor and antimicrobial resistance of *C. sakazakii*

Compared with the WT strain, the Δ*sdiA* mutant strain in normal culture showed significant differences in the expression of 244 genes, including 50 genes upregulated and 194 genes downregulated (Fig. 5). The Δ*sdiA* mutant strain showed significant expression differences in small molecule metabolism, polysaccharide anabolism, nucleotide glucose anabolism, lipopolysaccharide anabolism, galactose anabolism, GDP-mannose metabolism and carbohydrate anabolism, as well as response to virus and phage shock protein, as shown in Fig. 6A. As shown in Fig. 6B, the Δ*sdiA* mutant showed significant differences in lipid and sugar metabolic pathways, including sphingolipid metabolism, fructose and mannose metabolism, amino and nucleotide sugar metabolism, carbon metabolism, cytochrome P450 drug metabolism, retinol metabolism and pentose phosphate pathways (Table S3 and S4). These results indicate that the QS transcription regulator *SdiA* plays an important role in the cellular activities of *C. sakazakii*, and the mechanism of *SdiA* in each metabolic pathway of *C. sakazakii* needs to be revealed in the future.

**Fig. 5.**
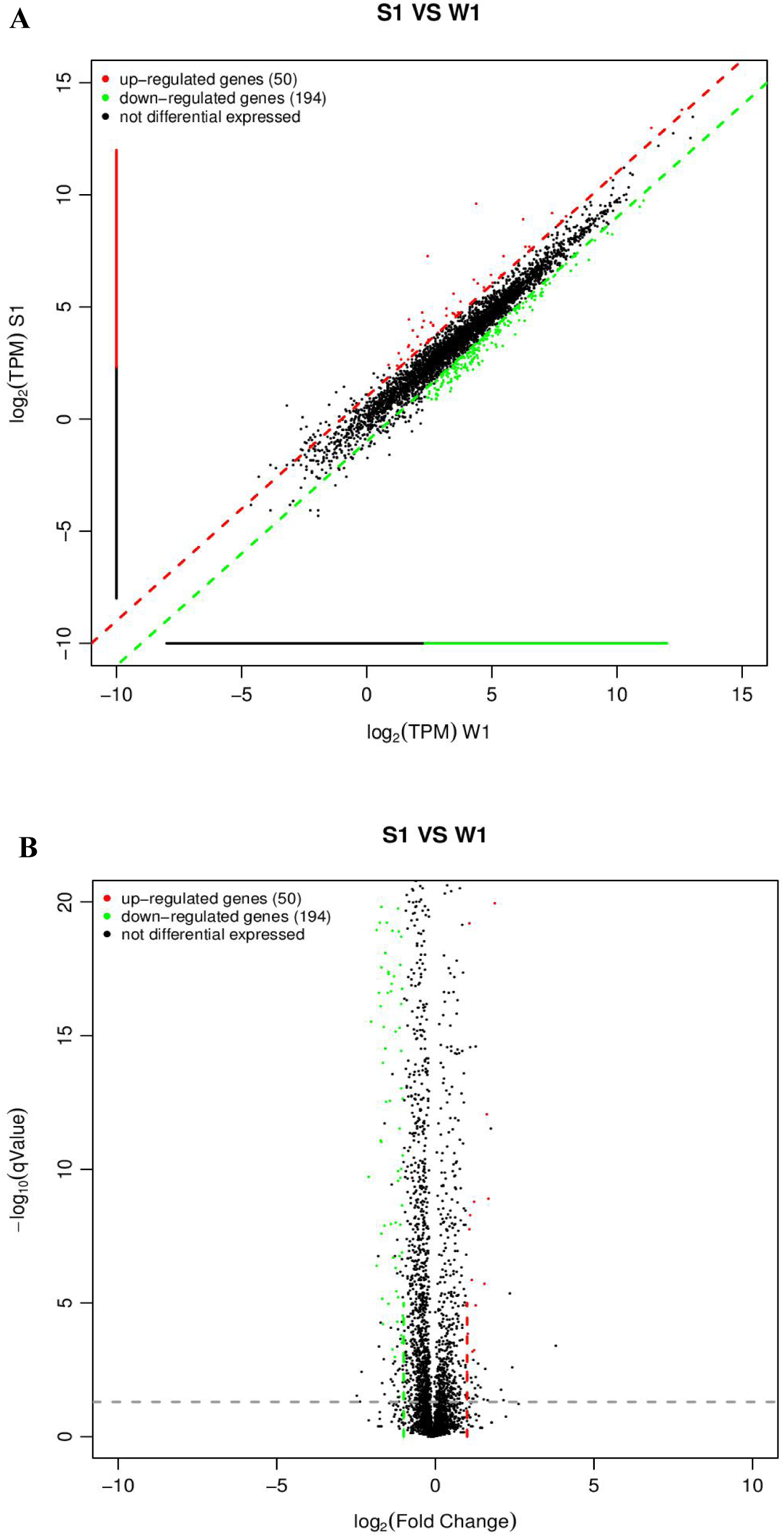
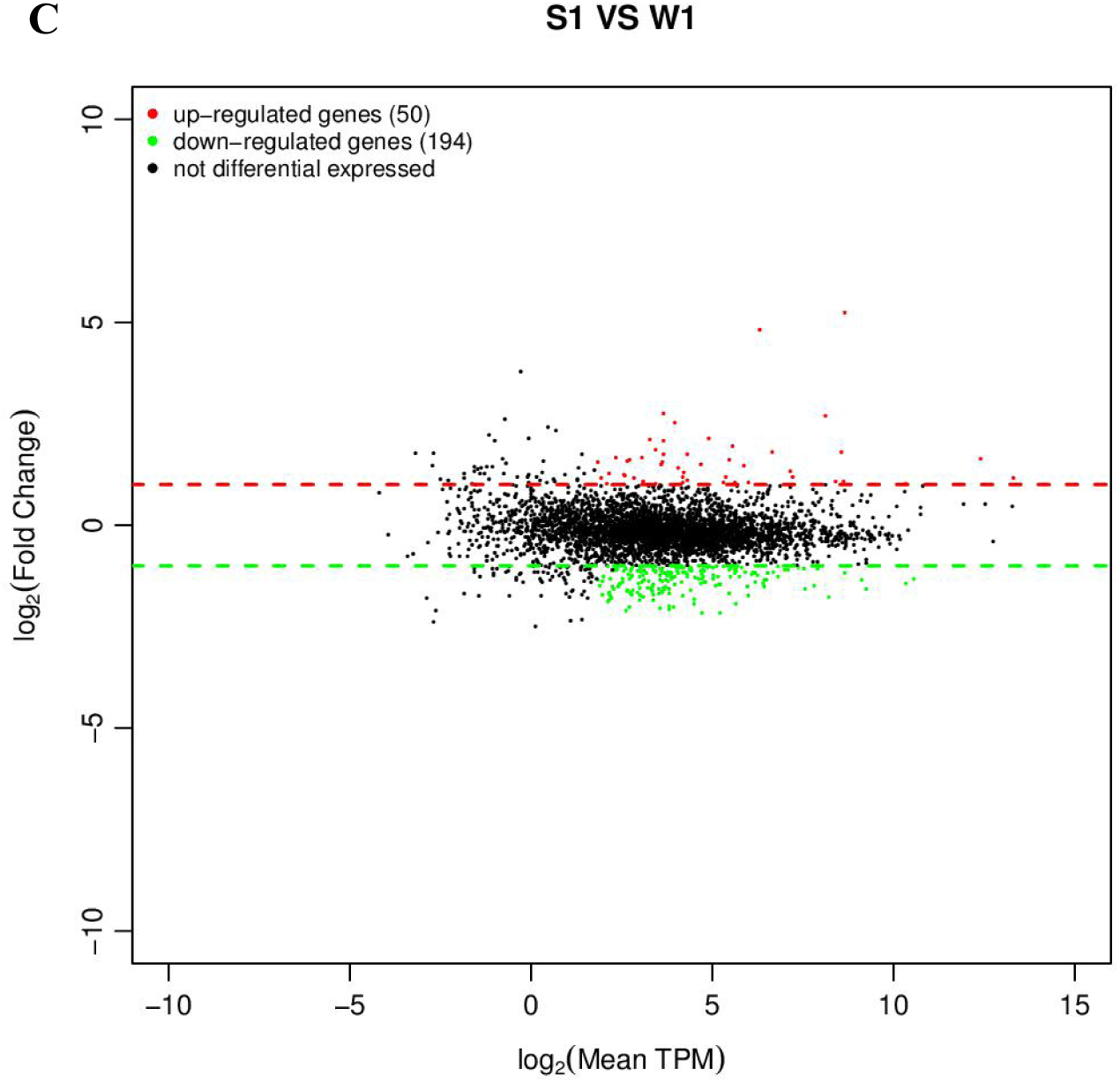
Expression difference analysis for transcriptional profiles (*C. sakazakii* WT (W1) and Δ*sdiA* mutant (S1) strains). **(A)** Comparison group expression difference scatter plot. The horizontal and vertical axes are respectively the log_2_ (TPM) values of two groups of samples. Each point in the figure represents a gene, and the closer the point expression is to the origin the lower. The vertical sample is relative to the horizontal sample. **(B)** Comparison group expression difference volcano diagram. The horizontal axis represents the fold-change (log_2_ (B/A)) value of gene expression difference between different groups of samples. The vertical axis represents gene expression pValue, the smaller the pValue, the greater the -log_10_ (pValue), and the more significant the difference. **(C)** MA diagram of expression difference between groups was compared. The horizontal axis is the mean value of log_2_ (TPM) of the two groups of samples ( (log_2_ (A) + log_2_ (B))/2). The vertical axis is log_2_ (Fold Change) (that is, log_2_ (B/A) ) values.

**Fig. 6.**
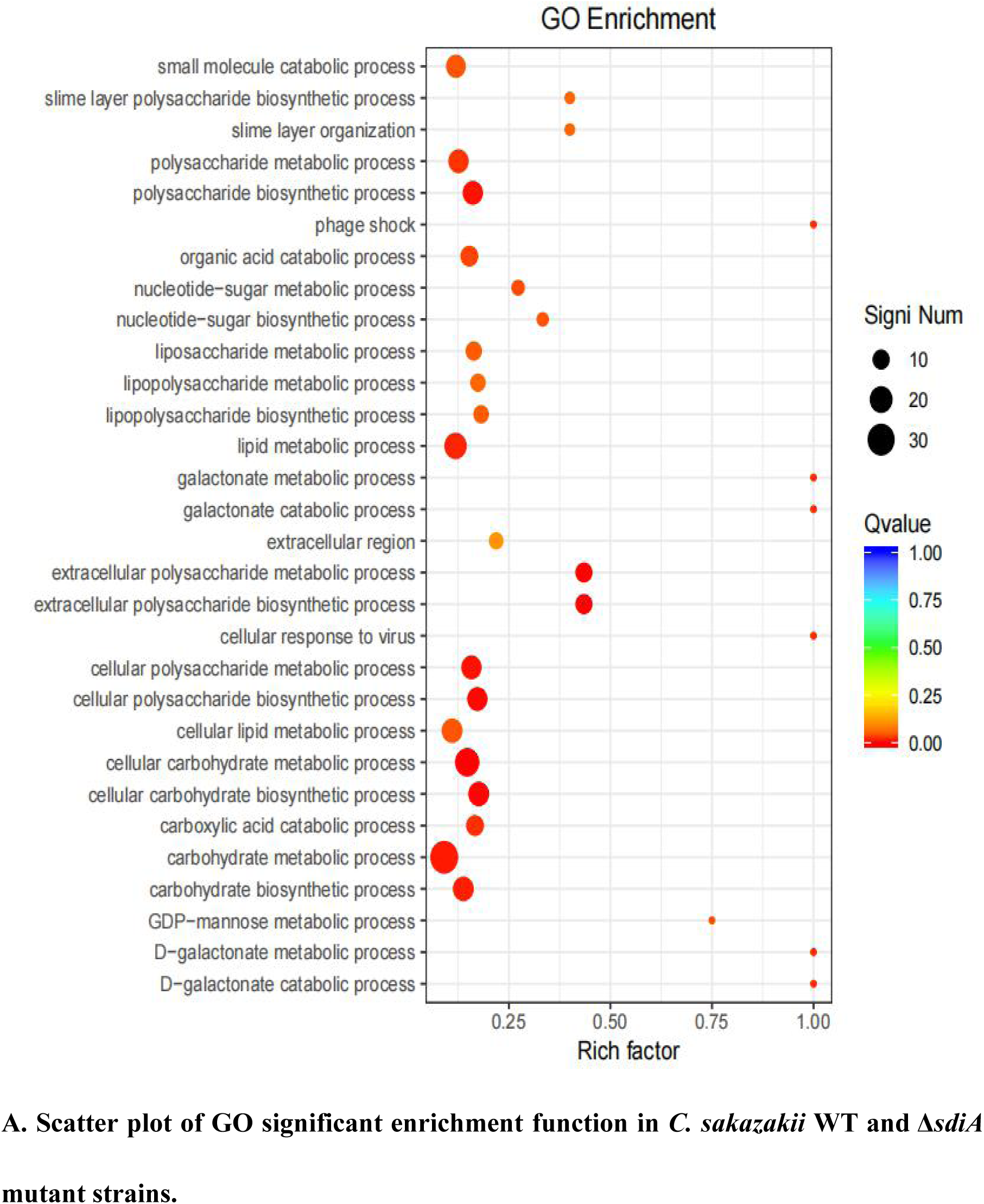
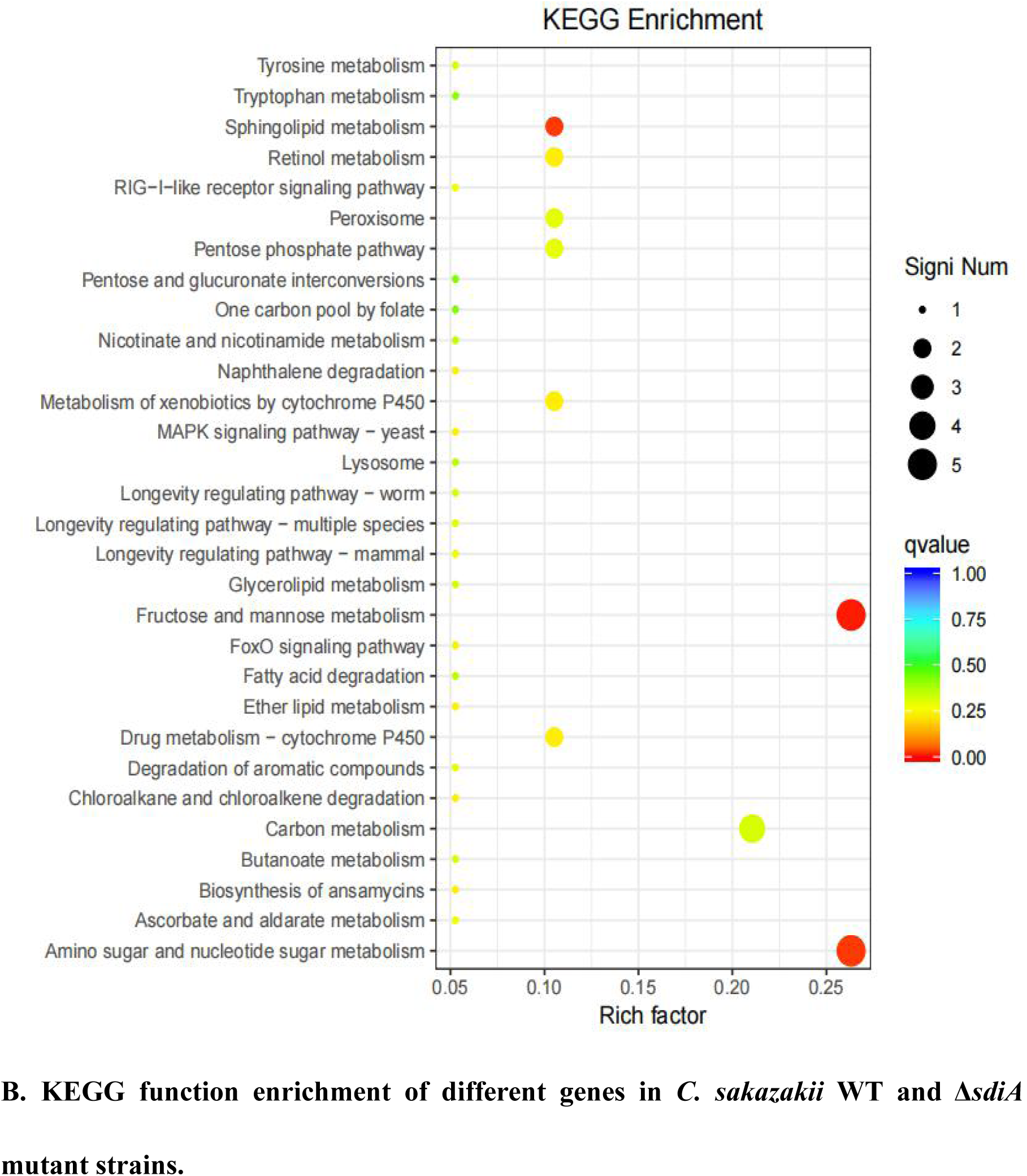
Genes enrichment analysis in *C. sakazakii* WT and Δ*sdiA* mutant strains. **(A)** Scatter plot of GO significant enrichment function in *C. sakazakii* WT and Δ*sdiA* mutant strains. **(B)** KEGG function enrichment of different genes in *C. sakazakii* WT and Δ*sdiA* mutant strains.

Compared to the WT strain, the expression of D-galactose operon-related genes in the Δ*sdiA* deletion strain was markedly upregulated, including the *dgoR*, *dgoK*, *dgoA*, *dgoD* and *dgoT* genes. In addition, the expression levels of the flagellate-related genes *FliA* and *FliC* were significantly increased, and the activity of flagella was enhanced. Consequently, the Δ*sdiA* mutant strain showed significantly enhanced motor activity. Among the upregulated genes, the 6-phospho-glucosidase gene *glvA* was the most significantly upregulated by 5.24-fold (Tables S1, S3 and S4). 6-Phosphoglucosidase has oxidoreductase activity, acting on CH-OH donors, NAD or NADP as receptors, and can bind to metal ions to express corresponding functions (31).

The differential gene expression of the *SdiA* gene deletion strain was markedly downregulated, which involved the VI secretion system and ABC transport system (21). The type VI secretion system is a multifunctional protein secretion system that can directly deliver toxins to eukaryotic cells and other bacteria, and its function is related to virulence host immune resistance and bacterial interactions (17). In the downregulated gene expression, the key function of the VI secretory system was the ATP-active membrane-associated protein VasK, whose gene expression was downregulated by 1.53-fold. The type VI secretion system plays an important role in the cell adhesion, virulence, invasion and proliferation of *C. sakazakii*; this system also produces protein toxin, which is a potential toxic factor (31). Moreover, the expression level of related genes in the ABC transport system was downregulated by 1.5-fold, which may affect the pathogenesis and virulence of *C. sakazakii* and reduce its ability to infect host cells and bacteria in drug resistance (Tables S2, S3 and S4)(29, 30).

## DISCUSSION

Homologous recombination is a conventional method for constructing gene deletion strains. Based on the principle of DNA homologous recombination, gene knockout technology can be roughly classified as homologous recombination, site-specific recombination and transposable recombination (32). QS is a process of communication between microbial cells, which is closely related to the secretion, colonization, invasion and infection of virulence factors of pathogenic bacteria and the formation of biofilms (14, 21). Nonetheless, the regulatory mechanism of QS on *C. sakazakii* remains largely unknown (25). To determine the role of QS in *C. sakazakii-*induced foodborne diseases, it is necessary to construct mutant strains of QS regulatory factors. Cao et al. successfully constructed *SdiA* frame deletion mutants using the suicide T vector pLP12 plasmid and *Escherichia coli* strain β2163 by sequence alignment based on the homologous recombination principle. Additionally, the QS receptor (*SdiA*) of *C. sakazakii* was screened out and the survival impact of *SdiA* on *C. sakazakii* under different environmental stresses was verified (25, 33). In this research, a mutant strain of the *C. sakazakii* QS receptor *SdiA* was successfully established by homologous recombination and electrical transformation. Moreover, the mechanism of *SdiA* action on the growth, drug resistance, motility, biofilm formation and adhesion of *C. sakazakii* was verified, which laid a foundation for *SdiA* targeted inhibition of the growth of *C. sakazakii* in PIF food.

Prediction of microbiology is a research field that applies mathematical models to predict the growth, inactivation and survival of microorganisms in food (34). The two-step method is a classical method to predict microbial modeling, but due to the two-step process, large errors will accumulate in the calculation (35). A one-step method is to construct the primary model and the secondary model simultaneously according to the observed values. Due to the one-step computation, the accumulated error is small, the fitting is excellent, and the growth parameters of microorganisms can be obtained more precisely (27). In recent years, the one-step predictive microbial model has been reported in the relevant literature, and a good fitting effect has been obtained. For example, the one-step kinetic analysis method was used to simultaneously analyze the growth curve of *Clostridium botulinum* in cooked beef (36) and the growth of *Salmonella* in liquid eggs (37). Fang et al. studied the growth kinetics of thermally damaged *C. sakazakii* in restored infant formula (RPIF) and established a mathematical model to forecast its growth (38). In RPIF, a logistic model was used to calculate the minimum and maximum growth temperatures of untreated cells as 6.5℃ and 51.4℃, respectively. The Huang model was used to calculate the minimum and maximum growth temperatures of *C. sakazakii* with heat treatment as 6.9℃ and 50.1℃, respectively. Although relevant literature has demonstrated the growth model of *C. sakazakii* in recovery PIF, the verification of the model by fluctuating temperature is deficient (39). The actual growth of microbes in food is based on practically a dynamic model. Therefore, we studied the actual growth of *C. Sakazakii* WT and Δ*sdiA* mutant strains in PIF based on one-step dynamic analysis. Pacheco et al. also found that no significant changes were observed in the growth curves of *Klebsiella pneumoniae SdiA* mutant and wild-type strains. However, cell division was impaired, and cell morphology was abnormal (19). This is similar to the results found in this study.

The increased motility of the *SdiA* mutant strain may be due to the influence of the *SdiA* gene on the expression of flagellate-related gene in *C. sakazakii*, which enhanced the motility of the mutant strain. Culler et al. studied the atypical *enteropathogenic Escherichia coli SdiA* gene and discovered that *SdiA* gene deletion demonstrated stronger motility (23). Kanamaru et al. showed that the flagellate-related gene *flic* of *Escherichia coli O157:H7* was negatively regulated by the *SdiA* gene (40), which was similar to the conclusion reached in this study. In addition, the cell morphology of the WT and Δ*sdiA* mutant strains was observed by scanning electron microscopy (Fig. 3B). When the *SdiA* gene of *C. Sakazakii* was removed, the overall morphology of the cells was not altered much. However, the Δ*sdiA* mutant strain was more elongated than the WT strain. This may help to enhance the swarming and swimming motility of *C. sakazakii*. Papenfort and Bassler showed that cyclic dimeric guanosine monophosphate (c-di-GMP) and cyclic adenosine monophosphate (cAMP), as QS signaling molecules, regulate bacterial virulence and biofilms. For example, the DSF-family autoinducer cis-2-dodecenoic acid (BDSF), an automatic inducer of the DSF family in *Bifidobacterium*, to *RpfR* leads to a decrease in intracellular c-di-GMP concentration, thereby affecting *Bifidobacterium* swarming motility, biofilm formation and virulence (21).

Culler et al. studied biofilm formation of atypical *enteropathogenic Escherichia coli* wild-type, *SdiA* mutant and complementary strains. The results showed that the *SdiA* mutant was able to form thicker biofilm structures and exhibit stronger motility. In addition, the transcription of *CsgA*, *csgD* and *FliC* was increased in the mutant strain, and the addition of AHLs reduced biofilm formation and the transcription of *csgD*, *CsgA* and *fimA* in the wild-type strain (23). Sabag et al. showed that AHL response genes and *SdiA-*dependent genes were identified in the *Enterobacter cloacae* mouse strain, and the presence of the *SdiA* gene inhibited the formation of biofilms (33). The conclusions from these studies are similar to those from this experiment, indicating that the *SdiA* gene may reduce the amount of bacterial biofilm formation. In this study, the expression levels of the *FliA* and *FliC* genes regulating the flagella of the *C. sakazakii SdiA* deletion strain increased at the transcriptional level, and the flagella activity was enhanced, which strengthened the adhesion process of the *SdiA* deletion strain in biofilm formation and was conducive to the formation of microcolonies. In a plant-related *Escherichia coli*, the *SdiA* mutant terminated the *csgBAC* operon, resulting in excessive pili production and increased adherence and biofilm formation of the *SdiA* mutant strain (41). Sharma and Bearson studied the effect of the QS transcriptional regulator *SdiA* on the retention of *Escherichia coli O157:H7* in weaned calves and found that *SdiA* could sense AHLs in the rumen of cattle and enhance the acid resistance of *Escherichia coli O157:H7* to allow *Escherichia coli O157:H7* to persist and colonize the bovine intestine for a period of time (22).

Stefany et al. showed that bacteria sense environmental changes by secreting their own inducers or receiving external environmental signals through the QS system to cope with the influence of bacterial population density, external pH and antibiotics, such as the antibiotic stress response (14). Tavio et al. studied the role of *SdiA* in multidrugresistant strains of *E. coli* and found that the expression of the *SdiA* gene promoted the expression of the multidrug resistance-related gene *acrAB*, leading to the strengthening of drug resistance in *E. coli*. In addition, amplification of *SdiA* can enhance resistance to *mitomycin C* by increasing the ability of chromosome replication and repair (41).

Carneiro et al. examined the expression of *SdiA* and genes associated with the nucleotide salvage pathway in *Salmonella* by qRT-PCR and found that *SdiA* affects glucose consumption, metabolism and gene expression in *Salmonella* and is closely related to metabolic pathways associated with glycine, purine, amino acid and aminoacyl tRNA biosynthesis. It plays a role in metabolic optimization under a high population density of *Salmonella* (42). Pacheco et al. found that *Klebsiella pneumoniae SdiA* binds to the promoter regions *imA*, LuxS, LSR-LSRA and *ftsQAZ*, and this binding is independent of AHLs. In addition, a lack of *SdiA* can increase the formation of *Klebsiella pneumoniae* biofilms and agglutination of yeast cells and lead to downregulation of type 3 and upregulation of type 1 pili expression. More importantly, *SdiA* inhibits bacterial adhesion and biofilm aggregation (14, 19, 21).

In conclusion, we constructed a mutant strain with the *SdiA* gene deletion of *C. sakazakii* CICC21550 using the principle of homologous recombination. The function of the QS-related gene *SdiA* in *C. sakazakii* was analyzed to obtain the relationship between the QS system and pathogenicity. The results demonstrated that *SdiA* enhanced the drug resistance of *C. sakazakii* but diminished its motility, adhesion and biofilm formation ability and had no effect on its growth. It can regulate the expression of D-galactose operon genes and flagellum-related gene upregulation and VI secretory system and ABC transport system-related gene expression downregulation. These results are helpful to further explore the function of the *SdiA* gene, revealing the pathogenic mechanism of *C. sakazakii*. Our data provide a new target for therapeutic interventions targeting the pathogenicity of *C. sakazakii* and developing quorum-sensing inhibitors.

## MATERIALS AND METHODS

### Bacterial strains and culture conditions

*C. sakazakii* CICC 21550 was obtained from the China Center of Industrial Culture Collection (CICC). *C. sakazakii* CICC 21550 wild-type (WT) and its mutant (Δ*sdiA*) strains were grown in trypticase soy broth (TSB) or on trypticase soy agar(TSA) at 37℃. Stock cultures of *C. sakazakii* CICC 21550 (WT) and mutant (Δ*sdiA*) strains were maintained in brain heart infusion (BHI) broth with 20% glycerol at -80℃. Activation of the strains was achieved by streaking the stock culture onto Violet red bile glucose agar (VRBGA) and incubating for 24 h at 37℃. The WT and Δ*sdiA* strains were stored on VRBGA at 4℃ to maintain cell viability.

### Sequence alignment, construction and identification of the *SdiA* deletion mutant

The nucleotide sequences of the *SdiA* gene and its upstream and downstream genes were obtained based on the whole genome sequencing results of *C. sakazakii* CICC 21550. Primers were designed using Primer 5.0 software (as shown in Table 1). All primers were synthesized by Sangon Biotech Co., Ltd. (Shanghai, China). *C. sakazakii* CICC 21550 genomic DNA was isolated using an Ezup Column Bacteria Genomic DNA Purification kit according to the manufacturer’s recommended protocol and stored at -20℃.

**Table 1.**
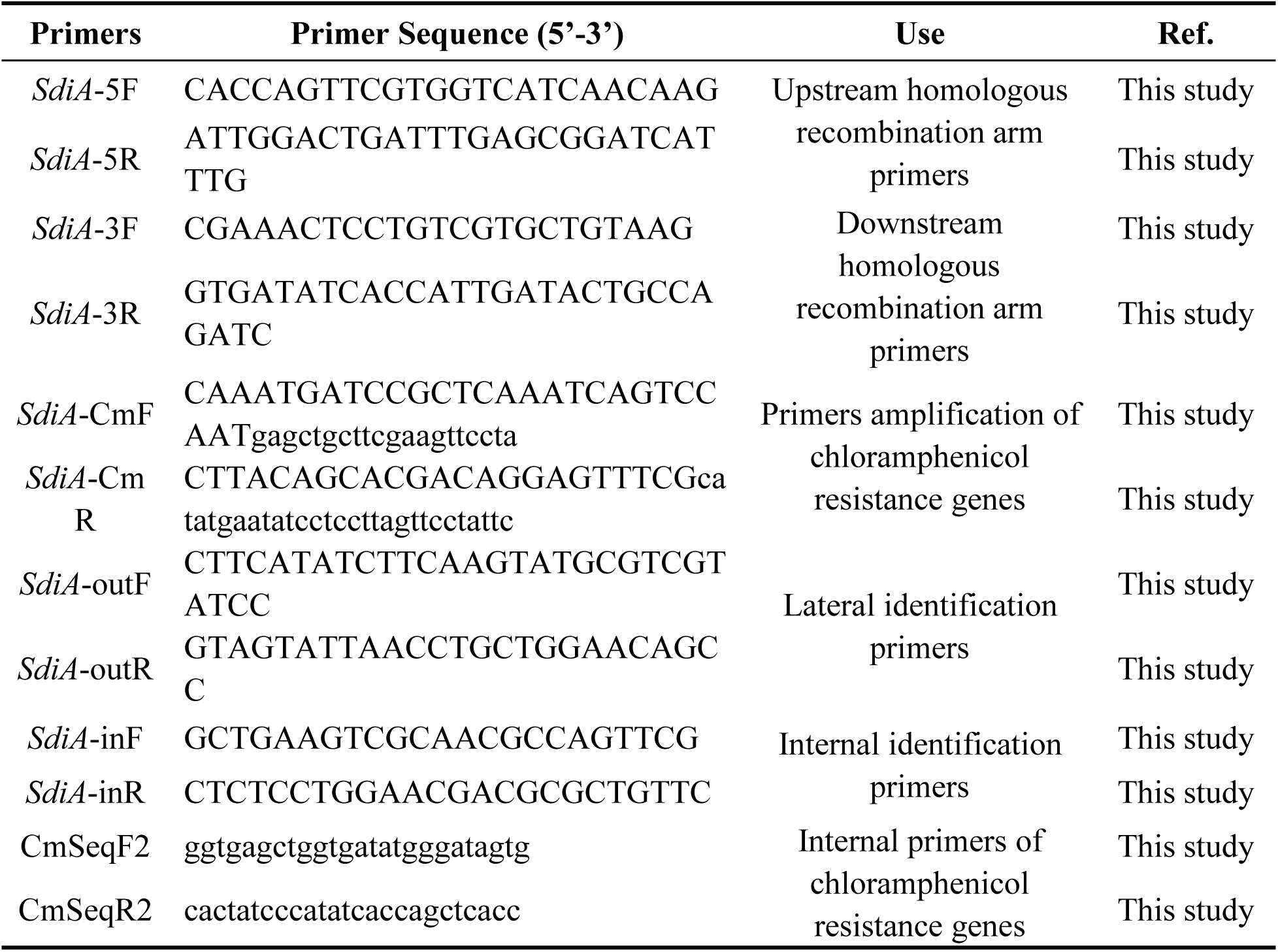
Primers used in this study.

The *SdiA* in-frame deletion mutant was constructed using the suicide plasmid pCVD442, plasmid pKD3 of the chloramphenicol resistance gene and *E. coli* strain β2155. These plasmids and *E. coli* strain β2155 were purchased from BioVector (Biovector NTCC Inc., China). The gene deletion construct was generated according to homologous recombination as previously described (22, 25, 32). Briefly, *SdiA* gene upstream and downstream homologous recombination arms were amplified by PCR from *C. sakazakii* CICC 21550 genomic DNA using primers *SdiA*-5F/*SdiA*-5R and *SdiA*-3F/*SdiA*-3R (Table 1). The Cm resistance cassette from plasmid pKD3 was PCR amplified with primers CmSeqF2 and CmSeqR2 (Table 1). The fusion PCR approach was used to construct the *SdiA* gene-targeting vector Δ*SdiA*::Cm (2650 bp, Fig. 1A), and the fusion PCR product was transformed into the suicide plasmid pCVD442. The plasmids were introduced by electroporation into *E. coli* strain β2155 and plated on TSA containing ampicillin (100 μg/ml) and 0.5 mM 2,6-diaminoheptanedioic acid (DAP) for selection at 37℃. After ligation, the pCVD442-Δ*SdiA*::Cm plasmid was transformed from β2155 into *C. sakazakii* CICC 21550 through conjugation (32). Finally, 100 μL was plated on TSA containing chloramphenicol (17 μg/ml) agar plates and cultured in an incubator at 30℃ until colonies formed.

The mutant strain was confirmed by PCR analysis using the Table 1 primer pair. Resistant colonies were streaked onto TSA plates without NaCl but supplemented with 10% sucrose (wt/vol) and chloramphenicol (17 μg/ml) to select for cells in which recombination and loss of the pCVD442 vector occurred. The chloramphenicol resistance gene was removed from the selected mutant using the pCP20 plasmid (Biovector NTCC Inc., China). Transformants were selected at 30°C on TSA plates containing 100 μg/ml ampicillin, and integration was confirmed by PCR with primers *SdiA*-outF/*SdiA*-outR (Table 1).

### Growth study

The powdered infant formula (PIF, 0-6 months, Yili Jinlingguanzhenhu) was sterilized by Co60 irradiation (8 kGy) and then sealed and preserved. A single colony of each strain from VRBGA was grown in 10 ml of TSB on a shaker at 130 rpm for 18-20 h at 37°C. Bacterial cells were harvested by centrifugation at 5000 rpm for 15 min at 4°C. Ten milliliters of 0.1% (wt/vol) peptone water (PW)-resuspended bacterial cells were washed off their surface toxins, and the supernatant was discarded. Then, the samples were resuspended in 10 ml of PW, and the bacterial washing step was repeated once (43). Subsequently, the solution was resuspended in 5 ml of PW and then diluted with PW to obtain an inoculum containing approximately 10^4.0-4.5^ CFU/ml or 4.0-4.5 log CFU/ml of *C. sakazakii*. According to the instructions of PIF, sterilized deionized water was used at high temperature and high pressure (121℃, 20 min) to prepare recovered milk powder. Then, 10 mL of milk was added to sterilizated empty test tubes with 0.1 mL aliquots of WT and Δ*sdiA* mutant cultures.

According to the growth parameters of *C. sakazakii* (38), the growth experiment with WT and Δ*sdiA* mutant strains was conducted at fluctuating temperatures. The inoculated samples were incubated at fluctuating temperatures of 8.4℃-48℃, 10℃-47.8℃, and 11℃-35℃. Samples were taken at predesignated time intervals and then serially diluted and plated on TSA plates overnight and enumerated.

### Scanning Electron Microscopy

Cells were grown and harvested by the above methods, resuspended in 2.5% glutaraldehyde fixative and fixed at 4℃ for 10 h. Subsequently, the cells were washed in 0.1 M phosphate buffered saline (PBS) and dehydrated through graded ethanol solutions. Before EM-30AX scanning electron microscopy (SEM, COXEM, Daejeon, Korea) observation, the samples were freeze-dried for 8 h and coated with gold.

### Swimming and swarming motility assays

The effect of WT and Δ*sdiA* mutant strains on swimming motility was assessed by examining swimming motility on TSA containing 0.3% (wt/vol) agar. *C. sakazakii* was cultivated in TSB overnight at 37°C in a shaking incubator at 130 rpm. Bacterial cells were diluted to 10^6^ CFU/ml after stab-inoculation into 0.3% (wt/vol) TSA agar plates and incubated for 6 h-24 h at 30°C. Motility was assessed quantitatively by measuring the circular swimming motion of the growing motile *C. sakazakii* every three hours. The effect of WT and Δ*sdiA* mutant strains on swarming motility was assessed by examining swarming motility on TSA containing 0.5% (wt/vol) agar. The same dilution (as mentioned above) was used to inoculate 0.5% (wt/vol) TSA agar plates and incubated for 6 h-36 h at 30°C. Motility was assessed quantitatively by measuring the circular swarming motion of the growing motile *C. sakazakii* every six hours.

### Crystal violet staining biofilm assay

The biofilm forming capacities of the WT and Δ*sdiA* mutant strains were assessed by a crystal violet staining assay (44) with slight modifications. Bacterial cells were cultivated in TSB overnight at 37°C until an optical density at 595 nm (OD_595_) of 0.5 or 10^9^ CFU/ml was reached. Thirty microliters of bacterial overnight culture was inoculated into 150 μL of TSB liquid broth in sterile 96-well plates and incubated for 24 h at 37°C. Then, 200 μL of 99% methanol (vol/vol) was added to each well for 15 min to fix the biofilms. After washing three times with 200 μL of 0.1 M PBS and dried in a hot air oven at 55°C for 1 h. The plates were stained with 200 μL of 1% crystal violet for 30 min, washed with 0.1 M PBS two times and then air dried at 37°C. Finally, the remaining crystal violet was resuspended in 200 μL of 95% ethanol and measured at 570 nm (OD_570_) using a microplate reader (SpectraMax®i3x, Molecular Devices, USA). In addition, 200 μL of TSB was used as a blank control, and each sample was set up with six replicates.

### Cell adhesion assay

The cell adhesion capacities were measured according to the method of Fabíola (45) with slight modifications. The bacterial cells were diluted with PW to 10^8^ CFU/mL. The 2 cm × 2 cm samples (made of glass, plastic, *silicone hydrogel* and stainless steel) were soaked in 10 ml of bacterial solution, incubated for 15 min at 30℃, and then washed three times with 10 ml of 0.01 M PBS to remove unbound cells. Then, 10 ml of 0.01 M sterile PBS was added for full oscillation to release the adhered cells into the PBS. The adhesion value was determined by counting viable cells on TSA plates.

### Tolerance and resistance assays

The susceptibility of the WT and Δ*sdiA* mutant strains to different antibiotics was evaluated by the disc diffusion method. The bacterial cells were diluted with PW to 10^8^ CFU/mL, and 0.1 mL of bacterial culture was plated onto a TSA plate. The antibiotic resistance was checked against 6 different antibiotics (200 µg/mL and 500 µg/mL): kanamycin, chloramphenicol, rifampicin, erythromycin, tetracycline hydrochloride and penicillin (Macklin Inc., Shanghai, China). Sterilized deionized water was used as a blank control. The sterilized circular filter paper (Φ=8 mm) was soaked in antibiotic solutions and deionized water for 10 min. Then, the filter papers were removed with sterile tweezers, affixed to a dried TSA plate and incubated for 24 h at 37 °C, and the diameter of the antibacterial coil was measured.

### RNA-Seq analysis

*C. Sakazakii* WT (W1) and Δ*sdiA* mutant strains (S1) were selected for RNA-Seq analysis. Total RNA was extracted by a UNIQ-10 column TRIzol total RNA extraction kit using cultures (130 rpm at 37°C) from the logarithmic growth stage. A Ribo-off rRNA Depletion Kit (Bacteria) was used to remove rRNA from total RNA samples. The cDNA libraries of RNA samples were constructed using the VAHTS™ Stranded mRNA-seq V2 Library Prep Kit for Illumina® (Vazyme Biotech Co., Ltd, Nanjing, China). The amplified cDNA library was visualized by 8% polyacrylamide gel electrophoresis (PAGE).

High-throughput sequencing was performed using the Illumina HISeq™ 2500 Platform. Illumina HISeq™ raw image data files were analyzed by CASAVA Base calling and transformed into sequenced reads. The resulting sequences were then referred to as Raw Data or Raw Reads, and the results were stored in FASTQ file format. Trimmomatic software was used for data processing, the original data quality value and other information were counted, and FastQC software was used for visual evaluation of the sequencing data quality of the samples (Fig. S1, Tables S5 and S6). Reference to the whole genome sequencing information of the *C. Sakazakii* CICC21550 strain for gene alignment analysis, gene expression level analysis and differentially expression analysis. ClusterProfiler software was used for functional enrichment analysis and the main biological functions of differential expressed genes were obtained by comparing the GO (Gene Ontology) database. The related biological system information was obtained by comparative analysis with the KEGG (Kyoto Encyclopedia of Genes and Genomes) database. All experiments were performed in triplicate (Fig. S2).

### Statistical analyses

All experimental results are presented as the mean ± standard deviation, and statistical analysis was performed with SPSS 17.0. The significance of the WT and Δ*sdiA* mutant strains was established through a t test using P values < 0.05.

## ACKNOWLEDGMENTS

This work was financially supported by the National Natural Science Foundation of China (NSFC 31601393, 31401597), the National and Science Foundation of Fujian Province (2018J01696), Fujian Agricultural and Forestry University (KXb16012A) and Fujian Education Department (JAT160147), 13th Five-year Plan on Fuzhou Marine Economic Innovation and Development Demonstration City Project.

We would like thank Yanhong Liu at the U.S. Department of Agriculture for critical review of the manuscript.

We declare no conflicts of interest.

Chuansong Cheng conceptualized the experiments, draft preparation and analyzed the data. Xiaotong Yan performed the experiments and collected the experimental data. Binxiong Liu supervised the experimentation and analyzed the data. Tao Jiang, Ziwen Zhou, Dongwei Zhang, Huayan Wang and Dengyuan Chen collected the data. Changcheng Li designed the experiment, analyzed data and revised the manuscript. Ting Fang designed the experiment, secured funding and revised the manuscript. All authors reviewed, revised, and approved the final manuscript.

## Supplemental material

**Fig. S1** *C. sakazakii* WT (W1) and Δ*sdiA* mutant (S1) strains sequencing saturation curve.

**Fig. S2** *C. sakazakii* WT (W1) and Δ*sdiA* mutant (S1) strains repeat correlation check scatter plots.

**Table S1** Upregulated genes annotation (*C. sakazakii* Δ*sdiA* mutant (S1) were compared with WT (W1) strains).

**Table S2** Downregulated genes annotation (*C. sakazakii* Δ*sdiA* mutant (S1) were compared with WT (W1) strains).

**Table S3** GO pathway enrichment analysis of significantly different genes (*C. sakazakii* Δ*sdiA* mutant (S1) were compared with WT (W1) strains).

**Table S4** KEGG pathway enrichment analysis of significantly different genes (*C. sakazakii* Δ*sdiA* mutant (S1) were compared with WT (W1) strains).

**Table S5** Quality and statistics of transcriptome library sequencing data.

**Table S6** Comparison and statistics of transcripts data.

